# Mitochondrial Electron Transport Chain Inhibition Suppresses LPS-Induced Inflammatory Responses via TREM1/STAT3 Pathway in BV2 Microglia

**DOI:** 10.1101/2019.12.25.888529

**Authors:** Cuiyan Zhou, Jie Zhang, Weihai Ying

## Abstract

Mitochondrial damage and neuroinflammation belong to two of the most important pathological factors in multiple neurological disorders. However, the effect of mitochondrial damage of microglia on microglial activation under pathological conditions has remained unclear. In our current study, we used BV2 microglia as a cellular model to determine the effects of mitochondrial electron transport chain (ETC) inhibitors on LPS-induced inflammatory responses of microglia. We found that all of the three mitochondrial ETC inhibitors, including rotenone, sodium azide and antimycin A, significantly inhibited LPS-induced inflammatory responses of the microglia, assessed by determinations of the protein or mRNA levels of IL-1β, IL-6, TNF-α, iNOS and COX2. Nuclear translocation of NF-κB p65 subunit does not appear to play an important role in the mitochondrial ETC inhibition-produced suppression of microglial activation. Instead, our study found that the mitochondrial ETC inhibitors significantly attenuated not only the LPS-induced increase in the TREM1 levels - an amplifier of inflammatory process, but also the LPS-induced increase in the ratio of phosphorylated STAT3 / STAT3. In summary, our study has suggested that mitochondrial ETC inhibition of microglia can lead to suppression of LPS-induced microglial activation, which may be mediated by the inhibitory effects of mitochondrial ETC inhibition on the LPS-induced increases in the level of TREM1 and the ratio of p-STAT3 / STAT3. These findings have provided valuable information for elucidating the relationships between mitochondrial damage and neuroinflammation in multiple neurological diseases.

## Introduction

Mitochondrial damage (1–3) and neuroinflammation (4, 5) belong to two of the most important pathological factors in multiple neurological diseases such as Parkinson’s disease, Alzheimer’s disease and brain ischemia. Mitochondria is the main source of cellular energy by coupling the oxidation of pyruvate and fatty acids to the production of adenosine triphosphate (ATP) by the mitochondrial ETC (6). The mitochondrial damage in the neurological disorders can produce major pathological alterations by causing both impaired energy metabolism and increased oxidative stress (7).

Neuroinflammation is a main host defense response to injury (4, 8). Numerous studies have indicated that neuroinflammation is a critical pathological factor in multiple neurological disorders (4, 5, 9), although neuroinflammation can also produce beneficial effects under certain conditions (10). Excessive activation of microglia - the major immune cells in the brain - leads to the production of multiple pro-inflammatory factors, which play a key role in neuroinflammation (11, 12). Since inhibition of the detrimental effects of neuroinflammation has been an important therapeutic strategy for multiple neurological diseases, it is of great significance to determine the key factors that modulate neuroinflammation in the brain under multiple pathological conditions.

The major goal of our current study was to determine the effects of mitochondrial ETC inhibition on neuroinflammation by using BV2 microglia - a widely used microglia model - as a cellular model. It is established that mitochondrial damage of such brain cell types as neurons can lead to necrosis, resulting in increased neuroinflammation. However, the effect of mitochondrial damage of microglia on microglial activation has remained unclear. Our current study has shown that mitochondrial ETC inhibition of microglia can profoundly suppress neuroinflammation in LPS-treated BV2 microglia, which could be mediated by the inhibitory effects of mitochondrial ETC inhibition on the LPS-induced increases in the levels of TREM1 and phosphorylated STAT3.

## Materials and Methods

### Reagents

Rotenone (45656), LPS (L2880) and NADH (N4505) were purchased from Sigma Aldrich (St Louis, MI, USA). Antimycin A (ab141904) was purchased from Abcam (Cambridge, USA). Sodium azide (DB0613) and primers were purchased from Sangon Biotech (Shanghai, China).

### Cell Cultures

BV2 microglial cells were a gift from the Institute of Neurology, Ruijin Hospital (Shanghai, China). The cells were plated into 24-well or 12-well culture plates in Dulbecco’s modified Eagle medium (HyClone, Logan, UT, United States) supplemented with 10% fetal bovine serum (Gibco, Carlsbad, CA, United States), 100 units/ml penicillin and 100 μg/ml streptomycin at 37 °C in a humidified incubator under 5% CO_2_.

### Intracellular Lactate Dehydrogenase (LDH) Assay

Cell death was determined by measuring the intracellular lactate dehydrogenase (LDH) activity of the cells. In brief, cells were lysed for 20 min in lysing buffer containing 0.04% Triton X-100, 2 mM HEPES and 0.01% bovine serum albumin (pH 7.5). Then 50 μL cell lysates were mixed with 150 μL reaction buffer containing 0.34 mM NADH and 2.5 mM sodium pyruvate (pH 7.5). The changes of the A_340nm_ of the samples were monitored over 90 s. Percentage of cell death was calculated by normalizing the LDH values of samples to the LDH values of the lysates of control.

### Nitric Oxide Assay

Nitric oxide (NO) production by BV2 microglia was assessed by determining the levels of nitrite accumulated in the culture media using Griess Reagent (Beyotime, Jiangsu, China). According to the manufacturer’s protocols, 50 µl of culture media of BV2 cells was incubated with 50 µl of Griess reagent I, which was then mixed with 50 µl of Griess reagent II. The absorbance at 540 nm of the samples was assessed by a microplate reader (Synergy 2; Biotek, Winooski, VT, United States). The nitrite levels of the samples were normalized to the protein concentrations, which were determined by the Bicinchoninic Acid (BCA) assay.

### Realtime Reverse Transcription-polymerase Chain Reaction (RT-PCR)

For RT-PCR assays, BV2 microglia (3 × 10^5^ cells/well) was plated into 24-well plate. Total RNA from BV2 cells was isolated by Rapid Extraction Kit for Total RNA from Blood (BioTeke Corporation, Beijing, China). Complementary DNA was obtained using Reverse Transcription Kit (Takara, Kyoto, Japan) according to the manufacturer’s protocol. cDNA was subjected to PCR for amplification. The primers used in this study as following: IL-1β forward, (5’-AAGGGCTGCFTTCCAAACCTTTGAC-3’) and reverse, (5’-ATACTGCCTGCCTGAAGCTCTTGT-3’), IL-6 forward, (5-TCCATCCAGTTGCCTTCTTG-3’) and reverse, (5’-AAGCCTCCGACTTGTGAAGTG-3’); TNF-α forward, (5’-CCCTCACACTCAGATCATCTTCT-3’) and reverse, (5’-GCTACGACGTGGGCTACAG-3’); TREM1 forward, (5’-CTGGTGGTGACCAAGG GTTC-3’) and reverse, (5’-CTTGGGTAGGGATCGGGTTG-3’), and GAPDH forward, (5’-CCTGCACCACCAACTGCTTA-3’) and reverse, (5’-GGCCATCCACAGTCTTCTGA-3’).

### Western Blot Assays

The cells were harvested and lysed in RIPA buffer (Millipore, Temecula, California, USA) containing 1% protease inhibitor cocktail (CWBio, Beijing, China), 1% phosphatase inhibitor cocktail (CWBio, Beijing, China) and 1 mM phenylmethanesulfonyl fluoride. Lysates were centrifuged at 12,000 rpm for 10 min at 4 °C. Nuclear protein samples were isolated using nuclear protein and cytoplasmic protein extraction kit (Beyotime, Jiangsu, China) according to the manufacturer’s protocol. After quantifications of the protein samples using BCA Protein Assay Kit (Pierce Biotechnology, Rockford, Illinois, USA), 30 μg of total protein was electrophoresed through a 10% or 15% sodium dodecyl sulfate-polyacrylamide gel and then transferred to 0.45-μm nitrocellulose membranes. The blots were incubated overnight at 4 °C with primary antibodies. The antibody dilutions were as follows: anti-iNOS antibody: 1:1000 dilution; anti-COX2 antibody: 1:800 dilution; anti-NF-κB antibody: 1:8000 dilution; anti-LaminA/C antibody: 1:8000 dilution; anti-TREM1 antibody: 1:2000 dilution; anti-Tubulin antibody: 1:3000 dilution (Abcam, Cambridge, USA) anti-p-AKT antibody: 1:2000 dilution; anti-p-STAT3 antibody: 1:2000 dilution; anti-p-p38 antibody: 1:2000 dilution; anti-p-ERK antibody: 1:2000 dilution; anti-p-JNK antibody: 1:2000 dilution; anti-AKT antibody: 1:2000 dilution; anti-STAT3 antibody: 1:2000 dilution; anti-p38 antibody: 1:2000 dilution; anti-JNK antibody: 1:2000 dilution; anti-ERK antibody: 1:2000 dilution; anti-GAPDH antibody: 1:1000 dilution. (Cell Signaling Technology, Danvers, MA) And then incubated with horse radish peroxidase-conjugated secondary antibody (1:3000 dilution, Epitomics, Hangzhou, China) in TBST containing 1% BSA at room temperature for 1 h. The intensities of the bands were quantified by densitometry using Gel-Pro Analyzer (Media Cybernetics, Silver Spring, Maryland, USA).

### Statistical Analysis

Data were presented as mean ± SEM and analyzed by one way analysis of variance (ANOVA) followed by Student-Newman-Keuls *post hoc* test. *P* values less than 0.05 were considered statistically significant.

## Results

### 1. Mitochondrial ETC inhibitors significantly suppressed LPS-induced increases in the pro-inflammatory factors in BV2 microglia

We used rotenone, sodium azide and antimycin A - three widely used mitochondrial ETC inhibitors (13, 14) - to produce inhibition of the mitochondrial ETC of BV2 microglia. The concentrations of rotenone, sodium azide and antimycin A used in our study were 100 nM, 2.5 mM, and 50 nM, respectively, since at these concentrations the mitochondrial ETC inhibitors did not significantly affect the survival of BV2 microglia (Supplementary Figures S1).

There are several hallmarks of microglial activation, including increased levels of NO and increased protein or mRNA levels of iNOS, COX2, IL-1β, IL-6 and TNF-α (15). Our study found that 100 nM rotenone, 2.5 mM sodium azide and 50 nM antimycin A significantly attenuated the LPS-induced increases in the NO level of the cells (Figure 1a). LPS treatment led to significant increases in the protein levels of iNOS and COX2 of the cells, which were also significantly attenuated by the mitochondrial ETC inhibitors (Figures 1b - 1d). We further found that the mitochondrial ETC inhibitors significantly suppressed the LPS-induced increases in the mRNA levels of IL-1β, IL-6 and TNF-α (Figures 2a - 2c).

**Figure 1.**
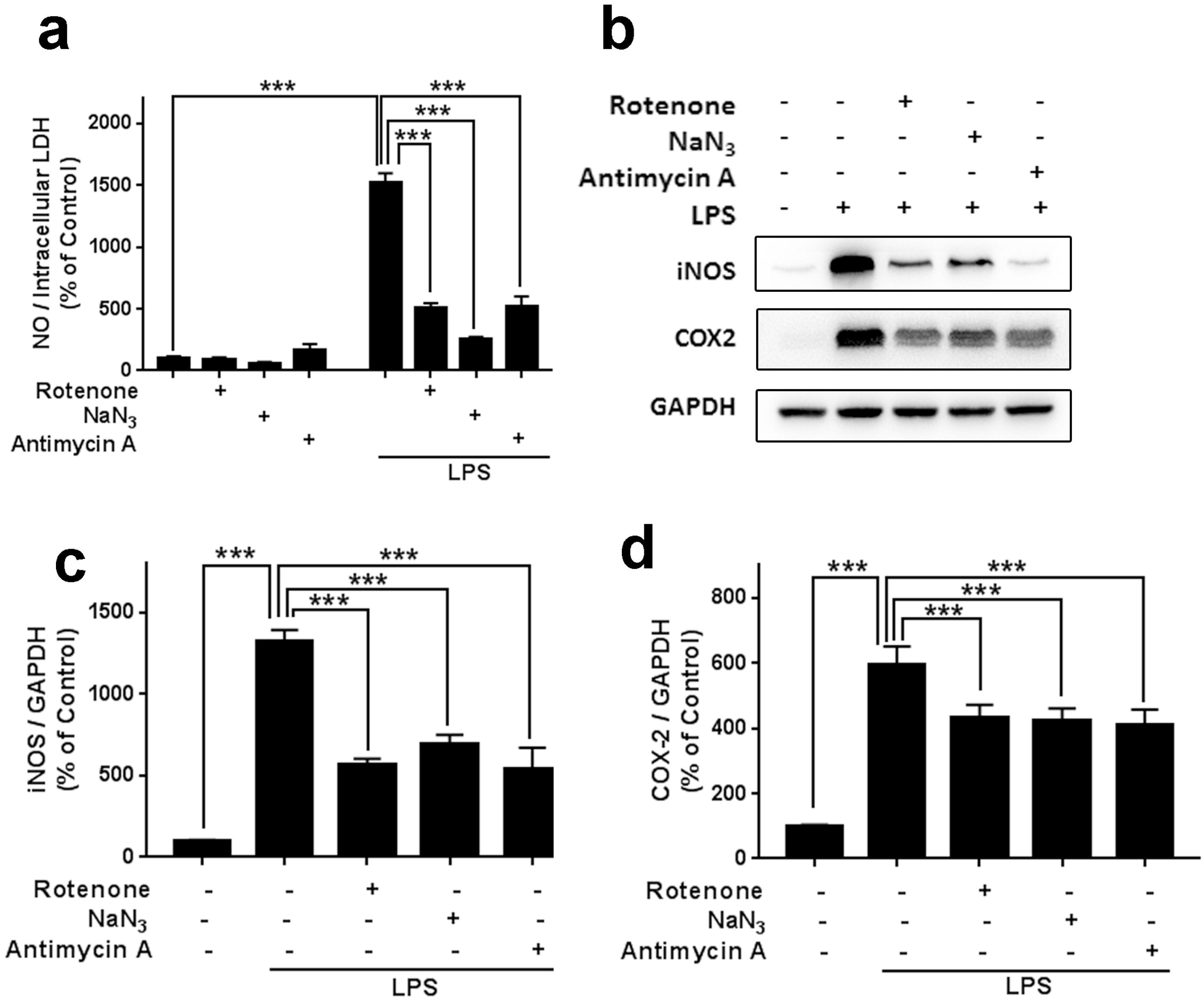
Mitochondrial ETC inhibitors significantly suppressed the increases in the LPS-induced inflammatory responses of BV2 microglia. (a) Rotenone, sodium azide and antimycin A significantly attenuated the LPS-induced increases in the NO level of the cells. (b) Western blot assays indicated that rotenone, sodium azide and antimycin A attenuated the LPS-induced increases in the protein levels of iNOS and COX2 of the cells. (c, d) Quantifications of the Western blot indicated that rotenone, sodium azide and antimycin A significantly attenuated the LPS-induced increases in the protein levels of iNOS and COX2 of the cells. After BV2 microglia were pre-treated with 100 nM rotenone, 2.5 mM sodium azide or 50 nM antimycin A for 30 min, the cells were incubated with 1 μg/ml LPS for 23.5 h. N = 12 - 15. The data were pooled from three independent experiments. ***, *P* < 0.001.

**Figure 2.**
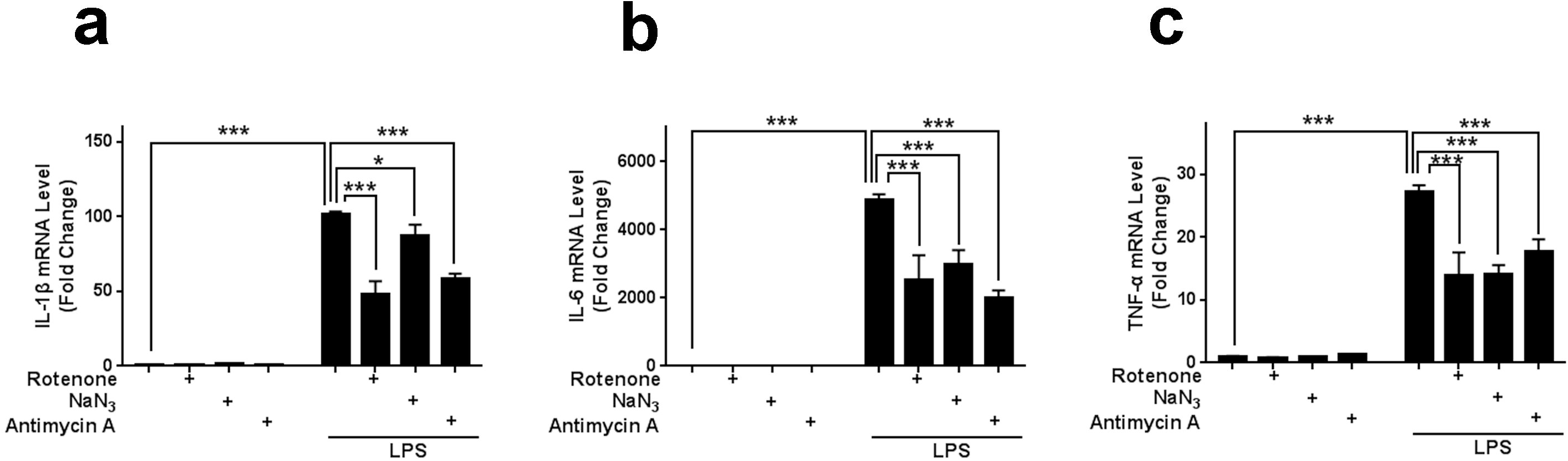
Mitochondrial ETC inhibitors significantly suppressed the LPS-induced increases in the mRNA levels of IL-1β, IL-6 and TNF-α in BV2 microglia. (a) Rotenone, sodium azide and antimycin A significantly attenuated the LPS-induced increase in the IL-1β mRNA level of the cells. (b) Rotenone, sodium azide and antimycin A significantly attenuated the LPS-induced increase in the IL-6 mRNA level of the cells. (c) Rotenone, sodium azide and antimycin A significantly attenuated the LPS-induced increases in the TNF-α mRNA level of the cells. After BV2 microglia were pre-treated with 100 nM rotenone, 2.5 mM sodium azide or 50 nM antimycin A for 30 min, the cells were incubated with 1 μg/ml LPS for 23.5 h. N = 6 - 8. The data were pooled from three independent experiments. *, *P* < 0.05; ***, *P* < 0.001.

### 2. The mitochondrial ETC inhibitors did not affect the nuclear translocation of NF-κB p65 subunit in the LPS-treated BV2 microglia

The nuclear factor NF-kB pathway is a well-established pro-inflammatory mediator (16), which plays critical roles in numerous inflammatory responses (17, 18). We conducted Western blot assays to determine the effects of the mitochondrial ETC inhibitors on the nuclear translocation of NF-κB p65 subunit - the major index of NF-κB activation (19), which did not indicate that the mitochondrial ETC inhibitors can significantly affect the nuclear translocation of NF-κB p65 subunit (Figures 3a – 3f).

**Figure 3.**
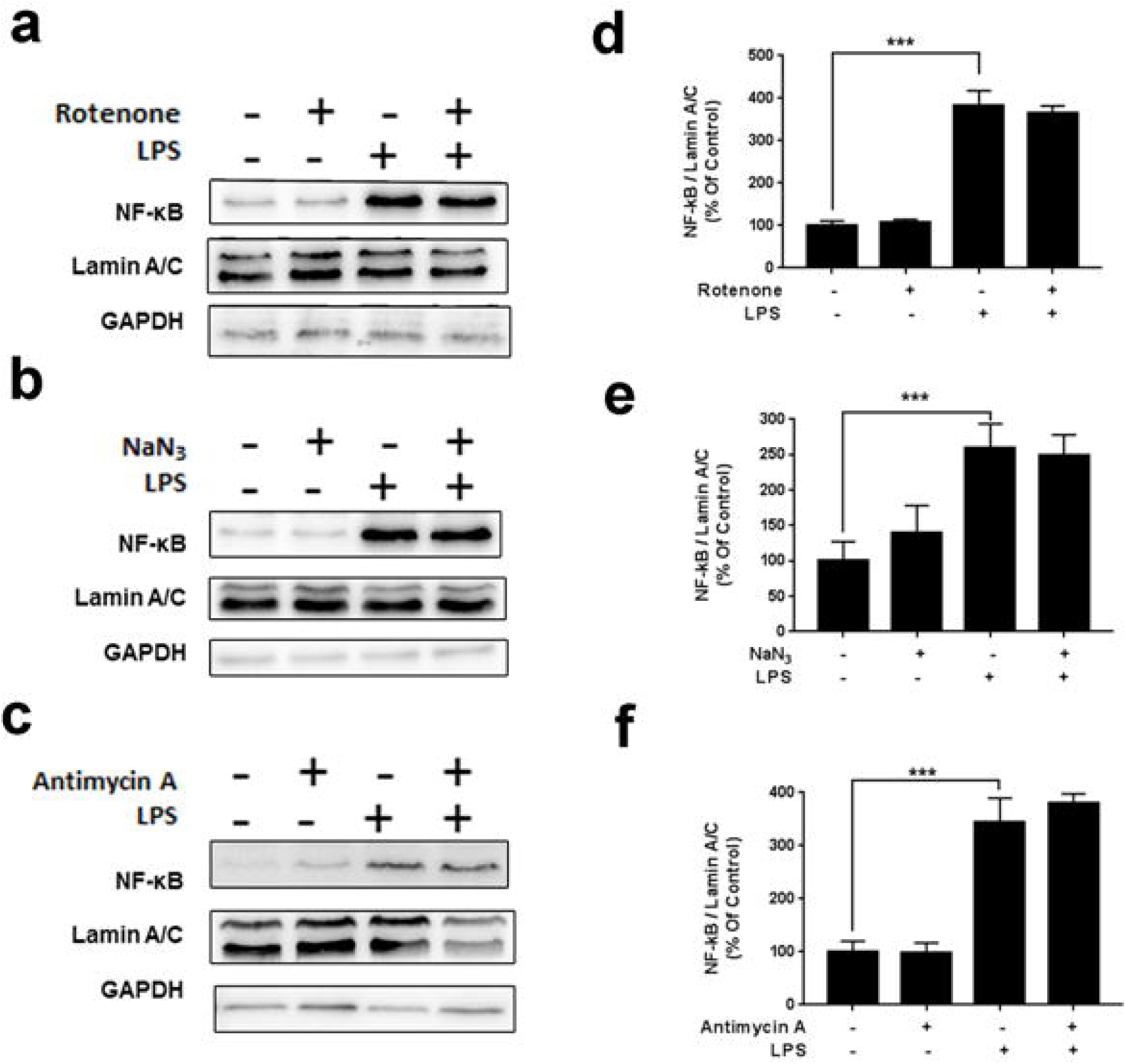
Mitochondrial ETC inhibitors did not significantly affect the nuclear translocation of NF-κB p65 subunit in LPS-treated BV2 microglia. (a, b, c) Western blot assays did not show that rotenone, sodium azide or antimycin A affected the nuclear translocation of NF-κB p65 subunit in LPS-treated BV2 microglia. (d, e, f) Quantifications of the Western blot did not show that rotenone, sodium azide or antimycin A significantly affected the nuclear translocation of NF-κB p65 subunit in LPS-treated BV2 microglia. N=7-9. The data were pooled from three independent experiments. ***, *P* < 0.001. After BV2 microglia were pre-treated with 100 nM rotenone, 2.5 mM sodium azide or 50 nM antimycin A for 30 min, the cells were incubated with 1 μg/ml LPS for 23.5 h. The images were representative of three independent experiments.

### 3. Three mitochondrial ETC inhibitors significantly suppressed the LPS-induced increases in both the mRNA and protein levels of TREM1 in BV2 microglia

A number of studies have suggested that TREMs play important regulatory roles in inflammation (20–22). We found that LPS treatment led to a significant increase in the mRNA level of TREM1, which was significantly attenuated by the three mitochondrial ETC inhibitors (Figures 4a). LPS treatment also led to a significant increase in the protein level of TREM1, which was attenuated by rotenone (Figure 4b), sodium azide (Figure 4c) and antimycin A (Figure 4d). Quantifications of the Western blot assays showed that the mitochondrial ETC inhibitors significantly suppressed the LPS-induced increase the protein levels of TREM1 in BV2 microglia (Figures 4e – 4g).

**Figure 4.**
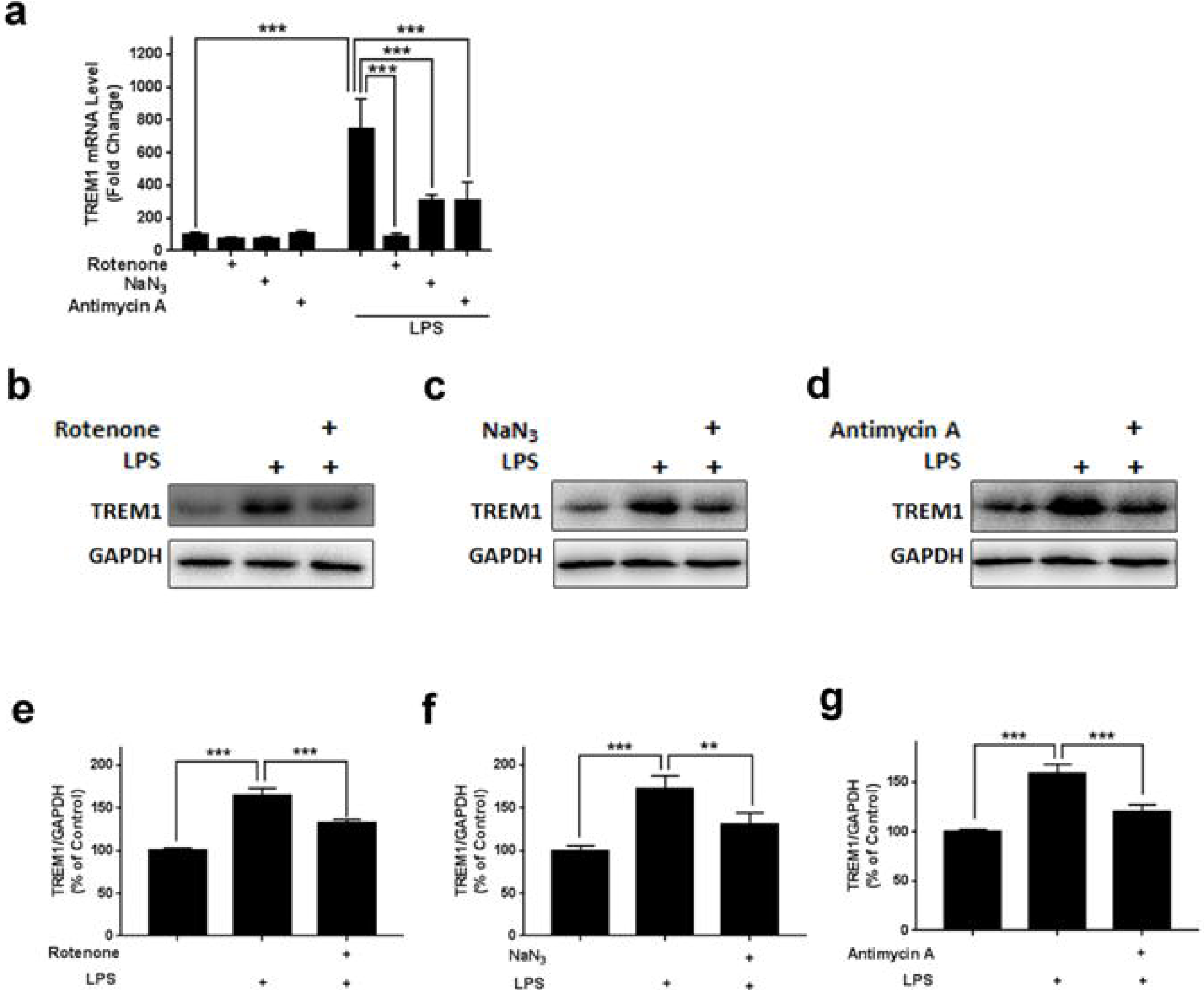
Mitochondrial ETC inhibitors significantly suppressed the LPS-induced increases in both the mRNA levels and the protein levels of TREM1 in BV2 microglia. (a) Rotenone, sodium azide and antimycin A significantly suppressed the LPS-induced increase in the TREM1 mRNA level in BV2 microglia. N = 7. The data were pooled from three independent experiments. ***, *P* < 0.001. (b, c, d) Western blot assays showed that rotenone, sodium azide and antimycin A attenuated the LPS-induced increase in the TREM1 protein level of the cells. (e, f, g) Quantifications of the Western blot assays indicated that all of the three mitochondrial ETC inhibitors significantly suppressed the LPS-induced increases in the protein level of TREM1 in BV2 microglia. After BV2 microglia were pre-treated with 100 nM rotenone, 2.5 mM sodium azide or 50 nM antimycin A for 30 min, the cells were incubated with 1 μg/ml LPS for 23.5 h. N = 12. The data were pooled from three independent experiments. **, *P* < 0.01; ***, *P* < 0.001.

### 4. Mitochondrial ETC inhibitors significantly suppressed the LPS-induced increase in the ratio of phosphorylated STAT3 (p-STAT3) / STAT3 in BV2 microglia

Since the key downstream proteins in the TREM1 signaling pathway include STAT3, MAPK, and AKT, we conducted Western blot assays to determine the effects of the three mitochondrial ETC inhibitors on the ratios of p-STAT3 / STAT3, p-p38 / p38, p-ERK1/2 / ERK1/2, p-JNK / JNK and p-AKT /AKT in the LPS-treated BV2 microglia (Fig. 5a). Of the five proteins, only the LPS-induced increase in the ratio of p-STAT3 / STAT3 was significantly suppressed by the mitochondrial ETC inhibitors (Figures 5b – 5f).

**Figure 5.**
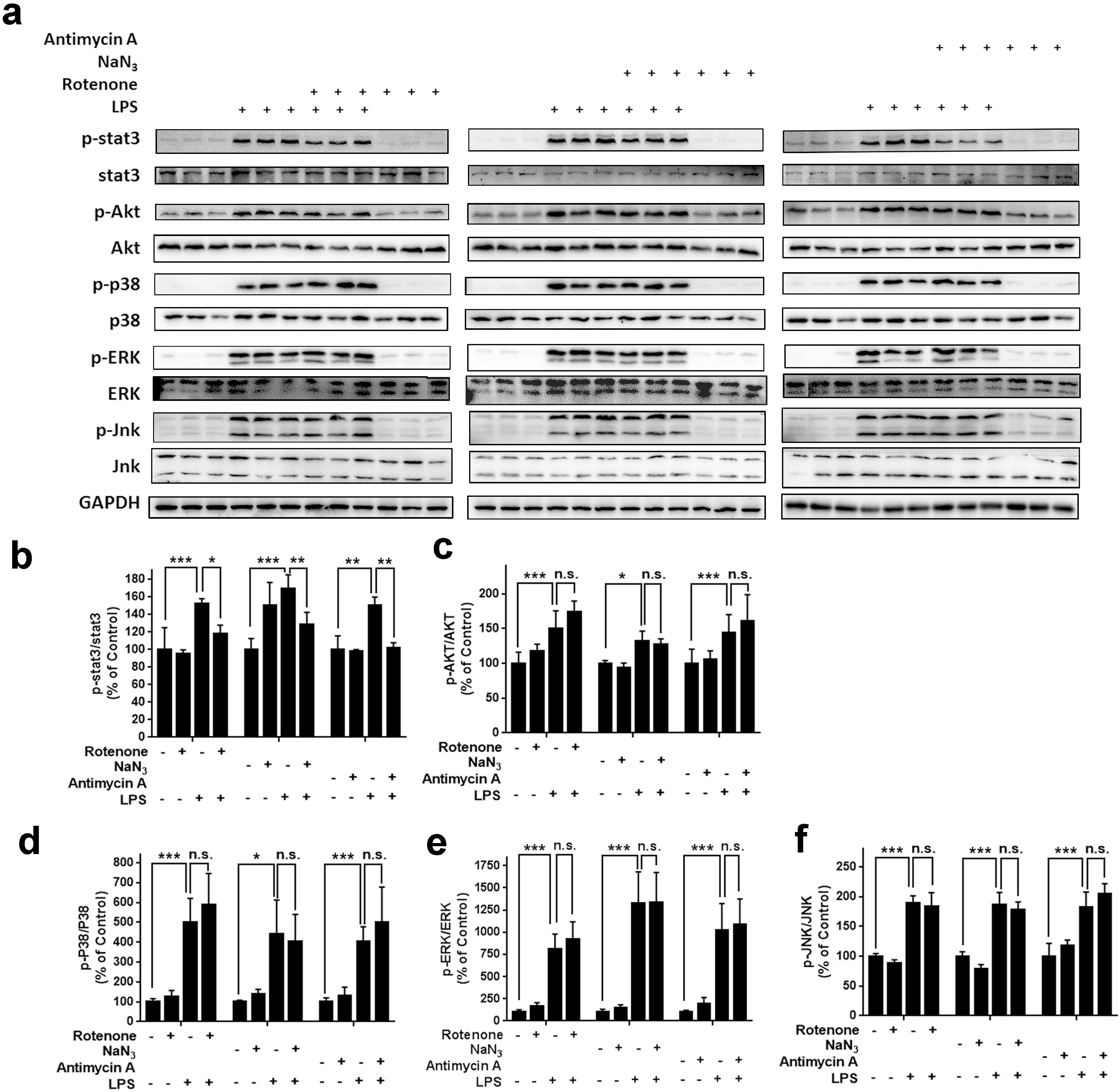
Mitochondrial ETC inhibitors significantly suppressed the LPS-induced increases in the ratio of p-STAT3 / STAT3 in BV2 microglia. (a) Western blot assays showed that rotenone, sodium azide and antimycin A suppressed the LPS-induced increases in the ratio of p-STAT3 / STAT3 of BV2 microglia. (b, c, d, e, f) Quantifications of the Western blot indicated that the three mitochondrial ETC inhibitors significantly decreased the ratio of p-STAT3 / STAT3, but not the ratios of p-p38 / p38, p-ERK1/2 / ERK1/2, p-JNK / JNK and p-AKT /AKT in the LPS-treated BV2 microglia. After BV2 microglia were pre-treated with 100 nM rotenone, 2.5 mM sodium azide or 50 nM antimycin A for 30 min, the cells were incubated with 1 μg/ml LPS for 30 min (for p-AKT, p-MAPK assays) or 2 h (for p-STAT3 assays). N = 6-15. The data were pooled from three independent experiments. *, *P* < 0.05; **, *P* < 0.01; ***, *P* < 0.001.

## Discussion

The major findings of our current study include: First, mitochondrial ETC inhibition can lead to decreased level of neuroinflammation in LPS-treated BV2 microglia; second, nuclear translocation of NF-κB p65 subunit does not play an important role in the mitochondrial ETC inhibition-produced suppression of microglial activation; third, mitochondrial ETC inhibition can decrease the LPS-induced increases in the levels of TREM1 in BV2 microglia; and fourth, mitochondrial ETC inhibition can suppress the LPS-induced increase in the ratio of p-STAT3 / STAT3 in BV2 microglia. These findings have provided novel information for elucidating the relationships between mitochondrial damage and neuroinflammation in multiple neurological disorders.

Mitochondrial damage and neuroinflammation belong to two most critical pathological factors in multiple neurological diseases. It is of significance to elucidate the relationship between these two pathological factors and expose the mechanisms underlying the relationship. However, the information regarding this relationship is highly insufficient. Our current study has shown that all of the three mitochondrial ETC inhibitors used in this study can significantly suppress the inflammatory responses in LPS-treated BV2 microglia. Our study has indicated that mitochondrial damage of microglia does not lead to increasing inflammatory responses in microglia. Instead, it leads to suppression of inflammatory responses in microglia. However, it is noteworthy that evaluation on the general effect of mitochondrial damage on neuroinflammation should be conducted on a broad scale: The mitochondrial damage of such cell types as neurons and astrocytes can lead to necrosis of the cells, which is expected to enhance the level of neuroinflammation.

It has been reported that uncoupling protein-2 (UCP-2) can provide neuroprotective effects by preventing neuronal death after stroke or brain trauma, while the mechanisms have remained unclear (23). Our study has provided a new line of information for accounting for the seemingly paradoxical finding: UCP-2 in microglia may lead to decreased microglial activation-based neuroinflammation by impairing microglial mitochondrial functions, resulting in the neuroprotective effect of UCP-2.

The transcriptional factors of NF-κB family play critical roles in inflammation and innate immunity (16, 24, 25). However, our study did not find that the mitochondrial ETC inhibitors can affect the nuclear translocation of NF-κB p65 subunit. Instead, our current study has indicated that mitochondrial ETC inhibition can decrease the LPS-induced increase in the level of TREM1 in BV2 microglia, which could at least partially account for the mitochondrial inhibitor-produced suppression of LPS-induced microglial activation. The triggering receptors expressed on myeloid cells (TREMs) are members of the immunoglobulin-like superfamily of receptors. In humans, two similar proteins, TREM1 and TREM2 are encoded by two homologous genes (26, 27). Accumulating evidence has indicated that TREM1 is an amplifier of the inflammatory process, while TREM2 has anti-inflammatory activity (22, 26, 28, 29). Inhibition of TREM1 has shown to attenuate the inflammation in several animal models of diseases, such as fatty liver diseases, rheumatoid arthritis or pneumonia (28, 30, 31). Our current study has shown that the three mitochondrial ETC inhibitors significantly decreased both the mRNA and protein levels of TREM1 in LPS-treated BV2 microglia, suggesting that the mitochondrial ETC inhibitor-produced suppression of the LPS-induced microglial activation results at least partially from its capacity to decrease the TREM1 level. However, it is warranted to further investigate the mechanisms underlying the capacity of the mitochondrial ETC inhibitors to decrease the TREM1 level.

We conducted studies to further determine if the mitochondrial ETC inhibitor can affect certain downstream factors in the TREM1-based signaling pathway. A number of studies have indicated that the key downstream factors in TREM1-based signaling pathway include Jak-STAT, MAPK, and AKT (32–34). Signal transducers and activators of transcription (STAT) plays critical roles in a variety of cellular functions including cell survival, proliferation, angiogenesis, invasion and immunity (35). Among the STAT family, phosphorylation of STAT3 is closely associated with inflammatory response (36). It was reported that TREM1 expression in splenic conventional dendritic cells was strongly up-regulated, which subsequently elicited the activation of STAT3 (37). Inhibition of STAT3 signal pathway was also shown to prevent α-synuclein-induced neuroinflammation and LPS-induced chronic liver injury (38, 39). Our current study has indicated that the mitochondrial ETC inhibitors can significantly decrease the ratio of p-STAT3 / STAT3 in the LPS-treated BV2 microglia. Collectively, these pieces of information has suggested that the mitochondrial ETC inhibitors lead to suppression of LPS-induced microglial activation by decreasing the ratio of p-STAT3 / STAT3 that results from the mitochondrial ETC inhibitor-produced decrease in the TRME1 level.

## Acknowledgment

The authors would like to acknowledge the funding supported by Shanghai Municipal Science and Technology Major Project (Grant No. 2017SHZDZX01) (to W.Y.).

## Legends of Supplemental Figure

**Supplemental Figure 1. Effects of rotenone, sodium azide and antimycin A treatment on the intracellular LDH levels of LPS-treated BV2 microglia.** Rotenone (a), sodium azide (b) and antimycin A (c) significantly decreased the intracellular LDH levels of BV2 microglia only at certain concentrations. After BV2 cells were pre-treated with rotenone (10, 100, 500 nM), sodium azide (0.25, 2.5, 10 mM) or antimycin A (10, 50, 100 nM) for 30 min, the cells were incubated with 1 μg/ml LPS for 23.5 h. N = 9. The data were pooled from three independent experiments. ***, *P* < 0.001.

## References

1. Arun S, Liu L, Donmez G. Mitochondrial Biology and Neurological Diseases. Current neuropharmacology. 2016;14(2):143–54.

2. Eid N, Ito Y, Horibe A, Otsuki Y, Kondo Y. Ethanol-Induced Mitochondrial Damage in Sertoli Cells is Associated with Parkin Overexpression and Activation of Mitophagy. Cells. 2019;8(3).

3. Golpich M, Amini E, Mohamed Z, Azman Ali R, Mohamed Ibrahim N, Ahmadiani A. Mitochondrial Dysfunction and Biogenesis in Neurodegenerative diseases: Pathogenesis and Treatment. CNS neuroscience & therapeutics. 2017;23(1):5–22.

4. Lucas SM, Rothwell NJ, Gibson RM. The role of inflammation in CNS injury and disease. Brit J Pharmacol. 2006;147:S232–S40.

5. Ward R, Zucca FA, Duyn JH, Crichton RR, Zecca L. The role of iron in brain ageing and neurodegenerative disorders. Lancet Neurol. 2014;13(10):1045–60.

6. Meyer A, Laverny G, Bernardi L, Charles AL, Alsaleh G, Pottecher J, et al. Mitochondria: An Organelle of Bacterial Origin Controlling Inflammation. Front Immunol. 2018;9.

7. Lin MT, Beal MF. Mitochondrial dysfunction and oxidative stress in neurodegenerative diseases. Nature. 2006;443(7113):787–95.

8. Becher B, Spath S, Goverman J. Cytokine networks in neuroinflammation. Nature reviews Immunology. 2017;17(1):49–59.

9. Bright F, Werry EL, Dobson-Stone C, Piguet O, Ittner LM, Halliday GM, et al. Neuroinflammation in frontotemporal dementia. Nature reviews Neurology. 2019;15(9):540–55.

10. Xanthos DN, Sandkuhler J. Neurogenic neuroinflammation: inflammatory CNS reactions in response to neuronal activity. Nature reviews Neuroscience. 2014;15(1):43–53.

11. Lloyd AF, Miron VE. The pro-remyelination properties of microglia in the central nervous system. Nature reviews Neurology. 2019;15(8):447–58.

12. Ferreira R, Bernardino L. Dual role of microglia in health and disease: pushing the balance toward repair. Frontiers in cellular neuroscience. 2015;9:51.

13. Han M, Im DS. Effects of mitochondrial inhibitors on cell viability in U937 monocytes under glucose deprivation. Arch Pharm Res. 2008;31(6):749–57.

14. Xiao F, Li Y, Luo L, Xie Y, Zeng M, Wang A, et al. Role of mitochondrial electron transport chain dysfunction in Cr(VI)-induced cytotoxicity in L-02 hepatocytes. Cellular physiology and biochemistry : international journal of experimental cellular physiology, biochemistry, and pharmacology. 2014;33(4):1013–25.

15. Coussens LM, Werb Z. Inflammation and cancer. Nature. 2002;420(6917):860–7.

16. Lawrence T. The nuclear factor NF-kappaB pathway in inflammation. Cold Spring Harbor perspectives in biology. 2009;1(6):a001651.

17. Tak PP, Firestein GS. NF-kappaB: a key role in inflammatory diseases. The Journal of clinical investigation. 2001;107(1):7–11.

18. Cao L, Li R, Chen X, Xue Y, Liu D. Neougonin A Inhibits Lipopolysaccharide-Induced Inflammatory Responses via Downregulation of the NF-kB Signaling Pathway in RAW 264.7 Macrophages. Inflammation. 2016;39(6):1939–48.

19. Hoesel B, Schmid JA. The complexity of NF-kappaB signaling in inflammation and cancer. Molecular cancer. 2013;12:86.

20. Bouchon A, Dietrich J, Colonna M. Cutting edge: Inflammatory responses can be triggered by TREM-1, a novel receptor expressed on neutrophils and monocytes. J Immunol. 2000;164(10):4991–5.

21. Bouchon A, Facchetti F, Weigand MA, Colonna M. TREM-1 amplifies inflammation and is a crucial mediator of septic shock. Nature. 2001;410(6832):1103–7.

22. Colonna M, Facchetti F. TREM-1 (triggering receptor expressed on myeloid cells): a new player in acute inflammatory responses. The Journal of infectious diseases. 2003;187 Suppl 2:S397–401.

23. Mattiasson G, Shamloo M, Gido G, Mathi K, Tomasevic G, Yi SL, et al. Uncoupling protein-2 prevents neuronal death and diminishes brain dysfunction after stroke and brain trauma. Nat Med. 2003;9(8):1062–8.

24. Lezoualc’h F, Behl C. Transcription factor NF-kappaB: friend or foe of neurons? Molecular psychiatry. 1998;3(1):15–20.

25. Mitchell JP, Carmody RJ. NF-kappa B and the Transcriptional Control of Inflammation. Int Rev Cel Mol Bio. 2018;335:41–84.

26. Walter J. The Triggering Receptor Expressed on Myeloid Cells 2: A Molecular Link of Neuroinflammation and Neurodegenerative Diseases. The Journal of biological chemistry. 2016;291(9):4334–41.

27. Tessarz AS, Cerwenka A. The TREM-1/DAP12 pathway. Immunology letters. 2008;116(2):111–6.

28. Rao S, Huang J, Shen Z, Xiang C, Zhang M, Lu X. Inhibition of TREM-1 attenuates inflammation and lipid accumulation in diet-induced nonalcoholic fatty liver disease. Journal of cellular biochemistry. 2019.

29. Ulrich JD, Holtzman DM. TREM2 Function in Alzheimer’s Disease and Neurodegeneration. Acs Chem Neurosci. 2016;7(4):420–7.

30. Fan DP, He XJ, Bian YQ, Guo QQ, Zheng K, Zhao YK, et al. Triptolide Modulates TREM-1 Signal Pathway to Inhibit the Inflammatory Response in Rheumatoid Arthritis. Int J Mol Sci. 2016;17(4).

31. Liu F, Zhang XG, Zhang B, Mao WW, Liu TT, Sun M, et al. TREM1: A positive regulator for inflammatory response via NF-kappa B pathway in A549 cells infected with Mycoplasma pneumoniae. Biomed Pharmacother. 2018;107:1466–72.

32. Fortin CF, Lesur O, Fulop T, Jr. Effects of TREM-1 activation in human neutrophils: activation of signaling pathways, recruitment into lipid rafts and association with TLR4. International immunology. 2007;19(1):41–50.

33. Arts RJW, Joosten LAB, van der Meer JWM, Netea MG. TREM-1: intracellular signaling pathways and interaction with pattern recognition receptors. J Leukocyte Biol. 2013;93(2):209–15.

34. Duan M, Wang ZC, Wang XY, Shi JY, Yang LX, Ding ZB, et al. TREM-1, an Inflammatory Modulator, is Expressed in Hepatocellular Carcinoma Cells and Significantly Promotes Tumor Progression. Ann Surg Oncol. 2015;22(9):3121–9.

35. Chong PSY, Chng WJ, de Mel S. STAT3: A Promising Therapeutic Target in Multiple Myeloma. Cancers. 2019;11(5).

36. Hillmer EJ, Zhang HY, Li HS, Watowich SS. STAT3 signaling in immunity. Cytokine Growth F R. 2016;31:1–15.

37. Gao S, Yuan LB, Wang YY, Hua CY. Enhanced expression of TREM-1 in splenic cDCs in lupus prone mice and it was modulated by miRNA-150. Mol Immunol. 2017;81:127–34.

38. Qin HW, Buckley JA, Li XR, Liu YD, Fox TH, Meares GP, et al. Inhibition of the JAK/STAT Pathway Protects Against alpha-Synuclein-Induced Neuroinflammation and Dopaminergic Neurodegeneration. J Neurosci. 2016;36(18):5144–59.

39. Yang Y, Zhao J, Song XQ, Li LF, Li FQ, Shang J, et al. Amygdalin reduces lipopolysaccharide-induced chronic liver injury in rats by down-regulating PI3K/AKT, JAK2/STAT3 and NF-kappa B signalling pathways. Artif Cell Nanomed B. 2019;47(1):2688–97.

