## Supplementary figures and images for "Mitochondrial Electron Transport Chain Inhibition Suppresses LPS-Induced Inflammatory Responses via TREM1/STAT3 Pathway in BV2 Microglia"

### Supplemental Fig. 1

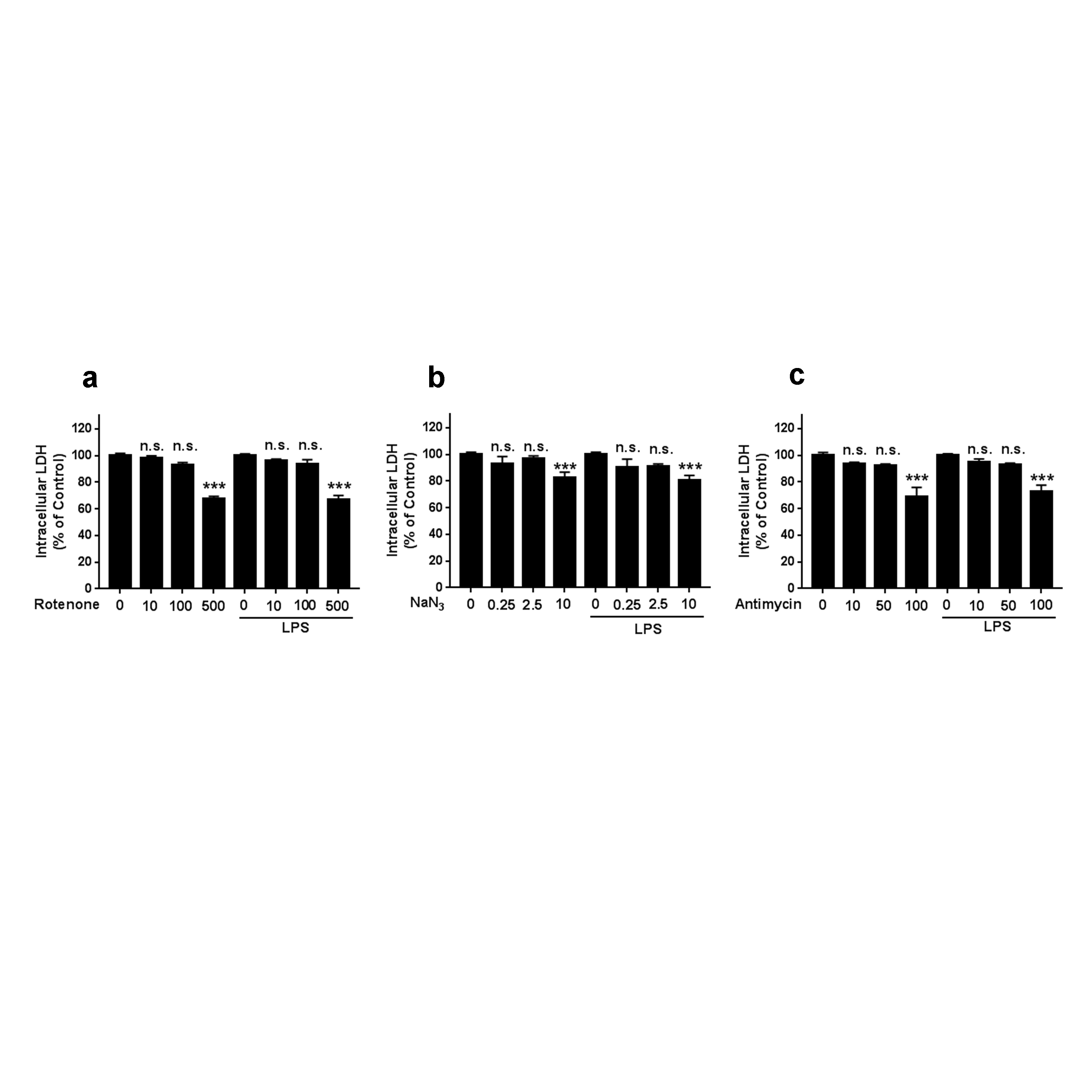
